# Genome Report: Chromosome-level draft assemblies of the snow leopard, African leopard, and tiger (*Panthera uncia, Panthera pardus pardus*, and *Panthera tigris*)

**DOI:** 10.1101/2022.04.26.489474

**Authors:** Ellie E. Armstrong, Michael G. Campana, Katherine A. Solari, Simon R. Morgan, Oliver A. Ryder, Vincent N. Naude, Gustaf Samelius, Koustubh Sharma, Elizabeth A. Hadly, Dmitri A. Petrov

**Author notes:** These authors contributed equally (Elizabeth A. Hadly, Dmitri A. Petrov).

## Abstract

The big cats (genus *Panthera*) represent some of the most popular and charismatic species on the planet. Although some reference genomes are available for this clade, few are at the chromosome level, inhibiting high-resolution genomic studies. Here, we assemble genomes from three members of the genus, the tiger (*Panthera tigris*), the snow leopard (*Panthera uncia*), and the African leopard (*Panthera pardus pardus*), at chromosome or near-chromosome level. We used a combination of short- and long-read technologies, as well as proximity ligation data from Hi-C technology, to achieve high continuity and contiguity for each individual. We hope these genomes will aid in further evolutionary and conservation research of this iconic group of mammals.

## Introduction

The genus *Panthera* comprises five extant species of Felidae and forms the majority of the group colloquially termed ‘big cats’. The clade is thought to have evolved approximately 5 million years ago [1,2] and its members are some of the most widespread and successful carnivores on the planet. However, over the last 100 years, all members of this clade have suffered widespread and severe declines, primarily due to anthropogenic causes [3]. As top terrestrial predators, members of the *Panthera* clade have naturally low abundances and slower intrinsic rates of population growth compared to prey species, forcing many populations into ‘threatened’ or ‘endangered’ listings by the IUCN and Convention on International Trade in Endangered Species (CITES), with severe ecological implications globally.

Due to their morphological similarities (not including their considerable variation in coat color and pattern), the genus *Panthera* has undergone substantial taxonomic rearrangements over the last century. Initially (1816-1916), the clade contained all spotted cats, irrespective of size or other morphological differences [4]. Later (1916), the clade was redefined to include only the jaguar (*P. onca*), the lion (*P. leo*), the leopard (*P. pardus*), and the tiger (*P. tigris*) [5]. At various times, arguments for the inclusion of the clouded leopard (*Neofelis nebulosa*) and the snow leopard (*P. uncia*) surfaced, but eventually genetic data supported the inclusion of only the snow leopard, defining the clade that we know today (jaguar, lion, leopard, tiger, snow leopard) [1,2,6]. Despite the refinement of the general tree topology, genomic data has been used to show that potential post-speciation hybridization and/or incomplete lineage sorting appears to have played a substantial role in the evolution of the genus [7–9]. However, when complex hybridization scenarios might explain a species’ history, or when events are recent enough that coalescent methods cannot be used, haplotype-level information such as is provided by contiguous, chromosome-level assemblies can greatly improve our understanding of these events.

To date, several draft genome assemblies have been published for members of *Panthera*, but few are at the chromosome-level, or contain many gaps. Draft genome assemblies have previously been published for the jaguar [7], lion [10], tiger [11–13], snow leopard [11], and leopard [14]. However, only assemblies for the lion [10] and the tiger (GCA_018350195.2) are highly contiguous and at the chromosome-level, so improved high-quality assemblies will facilitate both evolutionary research into the complex history and divergence patterns within the clade, as well as conservation applications, such as population genetics and forensic monitoring of wildlife products. There is no question that genomics can aid in the resolution of phylogenetic history, recently and notably identifying that giraffes (*Giraffa* spp.) are composed of several distinct lineages [15], and that the forest (*Loxodonta cyclotis*) and savannah elephant (*L. africana*) have been separated for 2.5 million years [16].

The tiger (*P. tigris*) comprises six extant subspecies, of which one (South China tiger, *P. t. amoyensis*) is functionally extinct in the wild and only exists in captivity [17]. Currently, draft reference genomes exist for the Bengal (*P. t. tigris*) [13,18], Amur (*P. t. altaica*) [11], and Malayan (*P. t. jacksoni*) [12], but not the Sumatran (*P. t. sumatrae*) and Indochinese (*P. t. corbetti*) tiger subspecies. Snow leopards (*P. uncia*), found in the high mountains of south-central Asia, are experiencing increasing threats and it is estimated that only about 3,500-7,500 of these cats remain in the wild [19–22]. At present, one fragmented genome assembly exists for the snow leopard, courtesy of DNAZoo (dnazoo.org). The leopard (*Panthera pardus*), the most widely distributed big cat, encompasses nine extant subspecies [6]. It is thought that African leopards (*P. p. pardus*) gave rise to eight Middle-Eastern (Arabian, *P. p. nimr*; Persian, *P. p. saxicolor*) and Asian (Indian, *P. p. fusca*; Sri Lankan, *P. p. kotiya;* Indochinese, *P. p. delacouri*; North-Chinese, *P. p. japonensis*; Amur, *P. p. orientalis*; and Javan *P. p. melas*) subspecies around 500–600 thousand years ago [23]. There are remarkably few genomic resources for leopards, with only a single fragmented *de novo* assembly from the Amur subspecies having been published to date [14].

Here, we add to the chromosome-level genome repertoire of *Panthera* with novel chromosome-level assemblies for the tiger, snow leopard, and African leopard. These resources should help guide further research into the evolutionary history and conservation of these big cats.

## Materials and Methods

### Tiger sequencing

The tiger sample used for whole genome assembly was provided by In-Sync Exotics (Wylie, TX, USA) from a male cub named “Kylo Ren’’ with unknown pedigree (Table 1). An EDTA whole-blood sample was taken during a routine veterinary check, subsequently packed on ice, and shipped to the Petrov and Hadly labs at Stanford University (Stanford, CA, USA). An aliquot of this sample was shipped on ice to Hudson Alpha (Huntsville, AL, USA) for 10x Genomics Chromium library preparation. This sample was extracted using the Qiagen MagAttract kit (Cat#:67563). A Chromium library was then prepared, shipped to Admera Health (Plainsfield, NJ, USA) and bidirectionally sequenced using 150bp reads on a lane of HiSeq X Ten (Illumina, Inc., San Diego, CA).

**Table 1:**
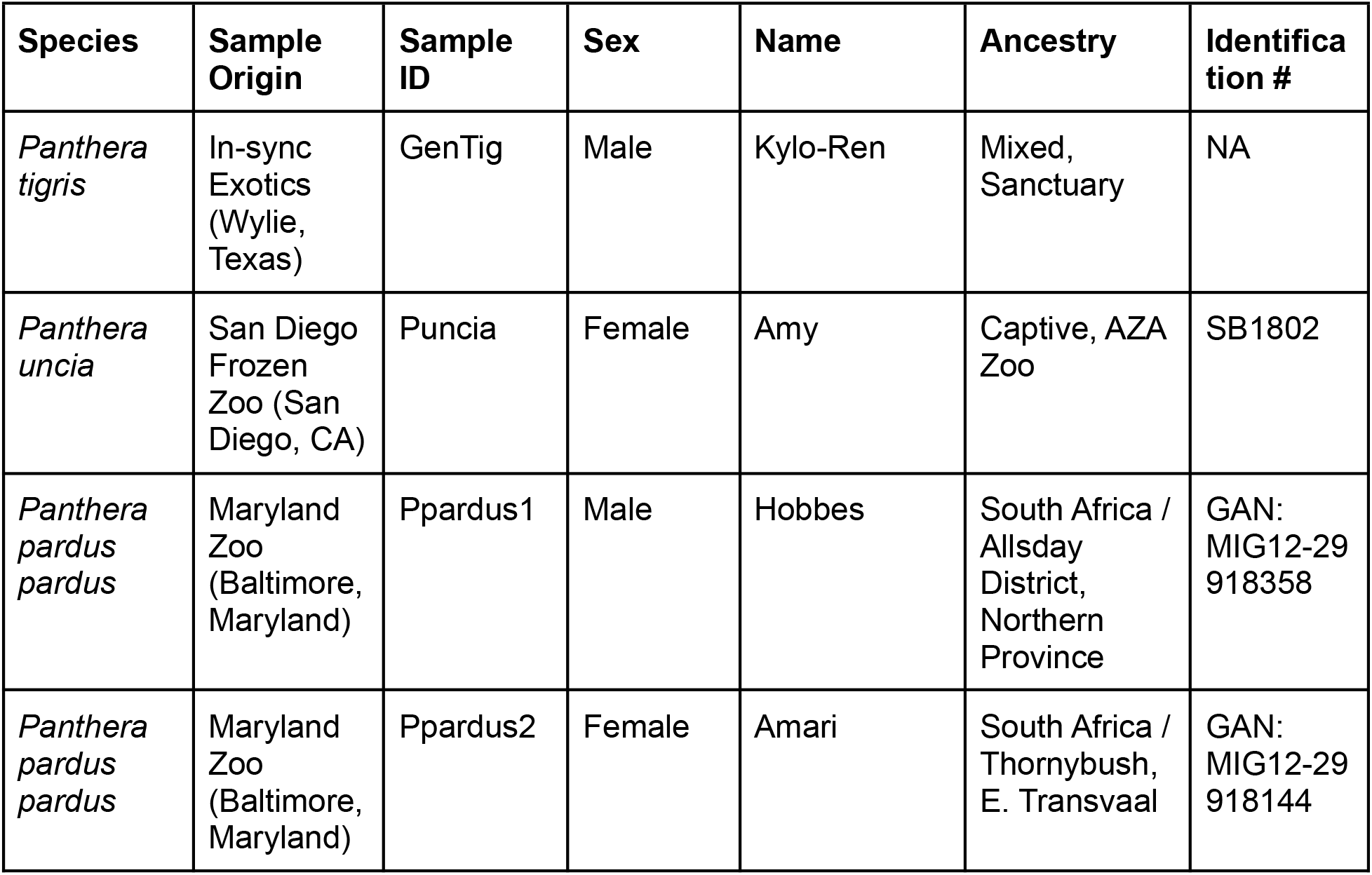
Sample information for individuals used in sequencing and *de novo* assembly

Material not used for 10x Genomics preparation was then used to construct a Hi-C library at Stanford University. Approximately 2mLs of whole blood was used as input in the Dovetail Hi-C kit (Cat#:21004) and a library was prepared according to the provided instructions. The finished library was then sent to Admera Health and bidirectionally sequenced using 150bp reads on a HiSeq X Ten lane.

### Snow leopard sequencing

An aliquot of a snow leopard cell line (ISIS#: 500107, Table 1) was obtained from the San Diego Frozen Zoo. The sample was sent to the Stanford Functional Genomics Facility (SFGF) for high molecular weight (HMW) DNA extraction using the Qiagen MagAttract kit (Cat#:67563) and 10x Genomics Chromium library preparation. The resulting library was shipped to Admera Health for sequencing on a Hiseq X Ten using bidirectional 150bp reads.

### African leopard sequencing

African leopard samples were obtained with permission from the Maryland Zoo (Baltimore, MD) from one male and one female leopard (Table 1). Samples consisted of previously biobanked EDTA whole-blood from individuals who are now deceased (ISIS#96040, 95007, Table 1). Whole blood was shipped on ice directly to Admera Health, where the samples were extracted using the Machery-Nagel NucleoBond HMW DNA extraction kit following the standard protocol. Samples were then shipped to the University of Georgia Genomics and Bioinformatics Core, where PacBio HiFi libraries were prepared and two Single Molecule, Real-Time (SMRT) cells were sequenced for each individual on the Sequel II. We note that these individuals were captured as cubs as the result of a potential poaching incident in South Africa, so the exact sample ancestry is unknown.

### Tiger genome assembly

The tiger genome was first assembled using 10x’s Supernova software (v2.1.1) using the default parameters. We output the assembly using the ‘--style=pseudohap’, ‘--minsize=10000’, and ‘--headers=sort’ flags in order to output a single, pseudohaploid assembly with short headers and scaffold sizes no less than 10,000bp. This genome was then used as the input assembly for the HiRise software from Dovetail Genomics. Briefly, the input *de novo* assembly and Dovetail Hi-C library reads were used as input data for HiRise, a software pipeline designed specifically for using proximity ligation data to scaffold genome assemblies [24]. Sequences from the Dovetail Hi-C library reads were aligned to the draft input assembly using a modified version of the SNAP read mapper (https://snap.cs.berkeley.edu). Separations of Dovetail Hi-C read pairs that were mapped to draft scaffolds were then analyzed by HiRise to produce a likelihood model for genomic distances between read pairs, and the model was then used to identify and break putative misjoins, score prospective joins, and to make joins above a defined threshold. Subsequent to scaffolding, shotgun sequences were used to close gaps between contigs. All Hi-C assembly steps were performed by Dovetail Genomics (Santa Cruz, CA), and the resulting assembly returned to us.

### Snow leopard genome assembly

The snow leopard genome was first assembled using the reads generated from the 10x Genomics Chromium library with the same methods outlined in the tiger genome assembly (see *Tiger genome assembly* above). Subsequently, Hi-C sequencing reads for the snow leopard generated by DNAZoo (dnazoo.org) were obtained and used with permission from the DNA Zoo Consortium (dnazoo.org) [25]. We used the juicer pipeline [26,27] to scaffold this assembly using the Hi-C sequence data. Briefly, the input genome assembly was first indexed with BWA index [28]. Subsequently, the generate_site_positions.py was used to generate positions of enzyme restriction sites of the input genome (the MboI enzyme was used as the restriction enzyme in this step). The juicer pipeline was then run with default settings using the draft assembly and previously generated enzyme restriction file as input. Finally, the 3D-DNA pipeline was used to generate a candidate assembly by inputting the 10x draft assembly again, along with the ‘merged_nodups.txt’ file generated in the previous step.

### Leopard genome assembly

We tested three assemblers (Flye; [29,30], wtdbg2; [31], and Hifi-asm; [32]) using the PacBio HiFi African leopard data. Using hifiasm0.9-r289, we ran standard commands with reads from both cells for each individual as input. We used flye2.8-b1674 standard commands (reads as input) with the ‘--genome-size’ flag set at 2.4g. Last, we used wtdbg2v2.5 with reads from both cells for each individual as input, the ‘-g’ flag set at 2.4g and the ‘-x’ flag set as ccs. Using the most complete assembly, as assessed by continuity statistics and gene completeness (see *Decontamination and assembly quality assessment* below), the primary Hifi-asm assemblies were then used as input into the juicer pipeline using the same pipeline as above. For this, we used Hi-C data generated by DNAzoo (DNAzoo.org) from the Amur leopard since contact mapping is unlikely to be impacted by the minimal divergence between these two subspecies [33].

### Decontamination and assembly quality assessment

Finalized draft genomes for each species were then uploaded to NCBI for contamination screening. The contamination file generated was then used to remove possible contaminants from each assembly by masking these regions with N’s using BEDtools2.26.0 maskfasta [34], or removed in the case of duplicated scaffolds. The cleaned genomes were then assessed for quality using Assemblathon2 [35] ‘assemblathon-stats’ scripts. We also assessed the continuity of the genome using BUSCO [36] with the mammalia_odb10 library, which evaluates the completeness of a set of manually curated orthologous genes.

We also calculated the depth of each library. Illumina reads were mapped to each genome using BWA-MEMv0.7.17 [37] and mean depth calculated using samtools depth [28] and awk. Briefly, each reference genome was indexed using bwa *index* with the flags ‘-a bwtsw’, reads mapped using bwa *mem*, and converted to bam files using samtools view with the flags ‘-bS’. For the genomes generated with Pacbio HiFi data, reads were mapped to each genome using minimap22-2.22 [38] to generate bam files and depth was calculated using the same method as for the Illumina libraries.

### Repeat masking and whole-genome alignment

The final decontaminated genome assembly for each individual was then repeat-masked using the same pipelines and settings as in [46]. Briefly, repeats were soft-masked first using RepeatMasker v4.0.9 based on known repeat databases from repbase using the flags ‘-species Carnivora’, as well as with flags ‘-nolow’, ‘-xsmall’, and ‘-gccalc’ [47,48]. Next, the masked genome produced in the previous step was used to build a database using RepeatModeler v1.0.11 BuildDatabase [47]. We then used RepeatModeler v1.0.11 to perform de novo repeat finding on the database produced in the previous step. Finally, a masked file with both known and de novo repeats was produced by running RepeatMasker v4.0.9 with the flags ‘-gccalc,’ and ‘-xsmall,’ and the library produced from the previous step as input with the initial masked file. The same steps were repeated to create a ‘hardmask’ of each genome for repeat analyses by simply replacing the ‘-xsmall’ and ‘-nolow’ flags with the ‘-a’ flag. Repeats were tabulated and plotted in R [39] with the ggplot2 package [40].

The resulting softmasked files were then used as input to LASTv921 [49] for whole-genome alignment. Each genome assembly was aligned to the autosomes and the X chromosomes from Felcat9 following scripts from https://github.com/mcfrith/last-genome-alignments. Genome alignment was visualized using the CIRCA software (http://omgenomics.com/circa) by plotting only the alignments with a mismap score of less than 1e-9 and the longest 50% from the major alignments for each of the Felcat9 autosomes and the X chromosome. The major alignment was determined as the scaffold belonging to the query assembly that represented a majority of the alignments for each of the Felcat9 autosomes and X chromosome.

After Hi-C integration, the snow leopard assembly was found to contain a misassembly which joined cat chromosomes E1 and F1. As a result, this scaffold was manually split using juicebox tools [26] and subsequently checked for accuracy by producing a realignment with LAST, but also through generation of an alignment dotplot with the program lastzv1.04.15 [41]. Alignments of both segments of the split scaffold and each of the E1 and F1 chromosomes were made in lastz using the flags ‘--notransition --step=20 --nogapped --format=rdotplot’ and resulting alignments were subsequently plotted in R (not shown). All previous stats (e.g. N50, BUSCO) were re-generated after the break was integrated.

### Leopard Ancestry Verification

In order to examine and verify the ancestry of the two putative African leopards, we used principal component analysis. Bam files generated from 37 African leopards [42] and bams that we generated for the two putative African leopards sequenced here (see *Decontamination and assembly quality assessment*), were input into angsdv0.931 [43] and PCAngsd [44] to generate a PCA. Angsd was first run using flags ‘-GL 2, -nThreads 64, -doGlf 2, -doMajorMinor 1, -SNP_pval 1e-6, and -doMaf 1’ to generate likelihood files for input into PCAngsd. PCAngsd was then run using flags ‘-e 3, -minMaf 0.05’ using the beagle files from the previous steps as input. PCA was visualized using R Core [39].

### Phylogenetics

Using the nucleic acid sequences for the recovered single-copy BUSCOs from each subspecies, we created a phylogenetic tree using only the unique subspecies of *Panthera*, and the clouded leopard (*Neofelis nebulosa*, DNAZoo.org; *Panthera uncia*, this study; *Panthera tigris altaica*, DNAZoo.org; *Panthera tigris jacksoni*, GCA_019348595.1; *Panthera onca*, DNAZoo.org; *Panthera leo*, GCA_018350215.1; *Panthera pardus pardus*, this study Ppardus1_PCG_1.0; *Panthera pardus orientalis*, DNAZoo.org). We did not include genomes that had mixed subspecies ancestry. Each gene was aligned using MAFFT v.7.490 [46] using the L-INS-i algorithm. Gene alignments were trimmed using Gblocks [51] v.091b. We inferred maximum likelihood individual gene trees using IQ-TREE 2 [52] v. 2.1.3 under default settings with 100 nonparametric bootstrap replicates and automated model selection using ModelFinder [53]. We concatenated the maximum likelihood trees as recommended by [54] and inferred the species tree using ASTRAL-III [48,49] v. 5.7.8 with 100 gene-only bootstrap replicates (flag ‘--gene-only’) and fully annotating the inferred tree (flag ‘-t 2’). For comparison, we also performed the same species-tree inference analysis using the concatenated consensus maximum-likelihood gene trees. The final species trees were rooted on *N. nebulosa*.

We also concatenated all Gblocks-trimmed gene alignments into a single unpartitioned concatenated alignment. We inferred maximum-likelihood trees with 100 bootstrap models using both reversible models using automated ModelFinder model selection (model GTR+F+R8 selected) and the non-reversible 12.12 model [55]. We then used FigTreev1.4.3 to visualize the tree (http://tree.bio.ed.ac.uk/software/figtree/). The final concatenated trees were rooted on *N. nebulosa*.

## Results and Discussion

### Sequencing and de novo genome assemblies

We generated sequencing for one 10x Genomics Chromium linked-read library for both a tiger and a snow leopard (Table 2). For the tiger, we generated a total of approximately 771 million paired reads (386 million read pairs), equating to an average genomic coverage of approximately 38× (Table 2). We additionally generated approximately 490 million paired reads from a Dovetail genomics Hi-C chromatin library, for an average genomic coverage of 29× (Table 2). For the snow leopard, we generated approximately 856 million paired reads for an average coverage of 40× (Table 2).

**Table 2:** Number of reads generated and average depth of sequencing for each African leopard. Depth was calculated by mapping circular consensus reads to the final assembly.

|  | **GenTig** | **Puncia** |
| --- | --- | --- |
| Number of 10x Chromium Linked Reads | 771,299,128 | 856,081,223 |
| Average Depth (10x) | 37.85× | 40.04× |
| Number of Hi-C Reads | 490,780,448 | NA |
| Average Depth (Hi-C) | 28.74× | NA |

Initial 10x assemblies for the tiger and the snow leopard generated 1,343 and 2,838 scaffolds, respectively. Both assemblies had similar contig N50 values (∼320kb; Table 3), however, the 10x assembly for the tiger had almost double the scaffold N50 and scaffold L90 of the snow leopard (Table 3). Interestingly, this does not appear to be a factor of the average input molecule size, which was 36,584 bases for the tiger and 36,914 bases for the snow leopard.

**Table 3:** Statistics of 10x Genomics Chromium assemblies for tiger and snow leopard.

|  | **Puncia (10x only)** | **GenTig (10x only)** |
| --- | --- | --- |
| Total length | 2,434,288,486 | 2,407,822,281 |
| Scaffolds | 2,838 | 1,343 |
| Contigs | 16,324 | 15,398 |
| Contig N50 | 322,039 | 328,032 |
| Scaffold N50 | 27,928,322 | 44,539,166 |
| Scaffold L90 | 96 | 66 |

However, the tiger did have a lower percent of duplicate sequences (8.73% in the tiger compared to 14.86% in the snow leopard), and also revealed a substantial difference in the number of molecules extending 10kb on both sides (111 in the tiger and 80 in the snow leopard). The snow leopard sample is additionally less heterozygous than the tiger, and the Supernova assembler estimated there to be 5.1kb between heterozygous SNPs in the snow leopard, compared to only 1.17kb in the tiger. While this would theoretically affect the assembler’s ability to phase the assemblies, it is not clear whether it would impact assembly quality. Indeed, the difference in coverage or molecule size alone does not appear to explain the differences between the assembly quality of the tiger and the snow leopard, since a previous study showed that 10x Genomics linked read data is robust across a range of coverages over 25× and is only greatly impacted by the mean molecule size [45].

Using PacBio HiFi sequencing, we generated data from two SMRT cells each for two African leopards of unknown origin. Of these, between ∼32–43% of the reads converted to circular consensus sequences, which translated to a final approximate depth of 25× and 27× for Ppardus1 and Ppardus2, respectively (CCS; Table 4). The low number of reads passing filter is likely due to the age and degradation of the samples, which were taken from each leopard post-mortem and stored in the Maryland Zoo biobank. Ppardus1 passed away in 2016 and Ppardus2 passed away in 2013.

**Table 4:** Number of reads generated and average depth of sequencing for each African leopard. ZMW: Zero-mode waveguides generated from SMRT sequencing.

|  | **Ppardus1 Cell 1** | **Ppardus1 Cell 2** | **Ppardus2 Cell 1** | **Ppardus2 Cell2** |
| --- | --- | --- | --- | --- |
| ZMWs generating circular consensus sequences | 1,908,091 | 1,695,169 | 1,893,243 | 2,001,644 |
| ZMWs filtered | 3,424,371 | 3,674,675 | 2,555,091 | 3,013,882 |
| Average depth | 25.06× |  | 26.98× |  |

Despite sample degradation and the resulting decrease in average coverage, data for both leopards were successfully used as input in three different genome assembly programs. In general, all assemblers produced draft assemblies of approximately 2.3-2.4Gb, but Hifi-asm [32] produced the assemblies with the fewest contigs (Ppardus1, 429; Ppardus2, 420; Table 5), and smallest contig L90 (Ppardus1, 108; Ppardus2, 94; Table 5). Comparatively, wtdbg2 [31] generated a draft assembly with 647 and 532 contigs for Ppardus1 and Ppardus2, respectively, while Flye [29], produced the largest number of draft scaffolds and the smallest contig N50s (Table 5). However, all long-read assemblers produced substantially more contiguous genomes than those produced by 10x Genomics linked-read technology, reiterating the utility of long-reads in genome assembly.

**Table 5:** Statistics comparing various pacbio assembly tools for Ppardus1 and Ppardus2. Assemblies chosen for Hi-C scaffolding are italicized.

|  | *HiFiasm*<br>*(Ppardus1)* | *HiFiasm*<br>*(Ppardus2)* | wtdbg2<br>(Ppardus1) | wtdbg2(Pp<br>ardus2) | Flye<br>(Ppardus1) | Flye<br>(Ppardus2) |
| --- | --- | --- | --- | --- | --- | --- |
| Total<br>length | 2,441,163,<br>814 | 2,436,970,3<br>22 | 2,377,391,<br>774 | 2,377,871,<br>889 | 2,416,305,<br>443 | 2,417,534,<br>344 |
| Contigs | 429 | 420 | 647 | 532 | 2,012 | 1,942 |
| Contig<br>N50 | 29,007,632 | 33,312,980 | 20,611,268 | 25,081,460 | 8,969,496 | 12,809,951 |
| Contig<br>L90 | 108 | 94 | 122 | 106 | 335 | 225 |

### Hi-C integration and comparison with other Panthera genomes

We lastly integrated Hi-C data for all four draft *de novo* assemblies produced in this study. In all cases, integrating Hi-C data led to an increase in continuity and a substantial decrease in the final L90 (Table 6). All assemblies except for the snow leopard have more than 90% of their genome contained in 38 or fewer scaffolds (the felid karyotype is 2n=38). This reinforces that these are all highly continuous, chromosome-scale assemblies.

**Table 6:** Assembly statistics of final assemblies for all species and individuals.

|  | **Ppardus1_PCG_1.0 (Pacbio + Hi-C)** | **Ppardus2_PCG_1.0 (Pacbio + Hi-C)** | **Puncia_PCG_1.0 (10x + Hi-C)** | **GenTig_PCG_1.0 (10x + Hi-C)** |
| --- | --- | --- | --- | --- |
| Total length | 2,441,326,314 | 2,437,106,822 | 2,434,048,986 | 2,407,461,690 |
| Scaffolds | 381 | 389 | 2,642 | 1,202 |
| Contigs | 706 | 674 | 16,601 | 15,399 |
| Contig N50 | 22,977,662 | 25,050,590 | 321,033 | 327,402 |
| Scaffold N50 | 126,810,544 | 158,060,271 | 112,063,229 | 145,225,440 |
| Scaffold L90 | 19 | 15 | 20 | 16 |

Compared to several of the previous tiger assembles that are publicly available (PanTig1.0, Cho et al., 2013 [7]; PanTig1.0-HiC, DNAzoo.org; Maltig1.0, [12]), GenTig_PCG_1.0 improved the contig N50 by almost four fold compared to PanTig1.0 and generated a comparable scaffold N50 to the improved PanTig1.0-HiC from DNAZoo and the ptimat1.1 assembly (Tables 6 and 7). However, the ptimat1.1 assembly, which was generated using long-read Pacbio sequencing and a trio-binning approach [46] has a larger contig N50 compared to all other assemblies. The newly generated snow leopard assembly (Puncia_PCG_1.0) showed a twofold increase in the Contig N50 (159.87kb to 321.03kb; Tables 6 and 7) and showed a similar scaffold N50 compared to the DNAZoo snow leopard assembly (PanUnc-HiC). The two African leopard assemblies showed an over 500-fold increase in the contig N50 compared to the published Amur leopard assemblies (PanPar1.0 and PanPar1.0_HiC; Tables 6 and 7). The African leopard genomes, which were assembled using PacBio HiFi data, also had higher contig N50s compared to the assemblies generated by 10x Genomics Chromium linked-read sequencing (Tables 6 and 7). Interestingly, the two leopard assemblies had slightly different contig N50 values which is likely explained by the number of reads that were able to be converted to circular consensus sequences (CCS; Table 4). Overall, long-read sequencing consistently improves the overall assembly continuity, especially impacting the distribution of contig lengths in the assembly.

**Table 7:** Summary statistics for other publicly available assemblies for the tiger, snow leopard, and leopard. Assemblies marked with * were used in phylogenetics analysis.

|  | **PanUnc-HiC**<br>(DNAZoo) | **PanPar1.0**<br>(GCA_001857705.1) | **PanPar1.0-HiC**<br>(DNAZoo) | **PanTig1.0**<br>(GCA_000464555.1) | **PanTig1.0-HiC**<br>(DNAZoo) | **Maltig1.0**<br>(GCA_019348595.1) | **ptimat1.1**<br>(GCA_018350195.2) |
| --- | --- | --- | --- | --- | --- | --- | --- |
| Total length | 2,408,534,935 | 2,578,019,207 | 2,578,248,701 | 2,391,383,019 | 2,391,065,193 | 2,424,644,036 | 2,408,695,688 |
| Scaffolds | 109,240 | 50,377 | 50,025 | 1,055 | 1,478 | 10,077 | 75 |
| Contigs | 159,866 | 162,036 | 162,190 | 126,631 | 126,369 | 36,281 | 140 |
| Contig N50 | 65,012 | 40,598 | 40,542 | 39,172 | 39,142 | 185,547 | 74,391,967 |
| Scaffold N50 | 141,103,905 | 21,701,857 | 155,751,443 | 8,860,407 | 145,958,240 | 21,297,815 | 146,942,463 |
| Scaffold L90 | 17 | 135 | 16 | 15 | 284 | 109 | 16 |

In order to further evaluate the quality of the assemblies, we ran BUSCO [36] on all available genomes for the leopard, tiger, and snow leopard. For the most part, all assemblies generated with linked-read or long read technology had minimal differences between BUSCOs found to be in single-copies, missing, fragmented or duplicated (Figure 1). Assemblies generated from both African leopards had a higher number of single-copy BUSCOs compared to either of the assemblies generated with the linked-read technology, except those generated with the wtdbg2 (Redbean; [31]) assembler (Figure 1). Overall, there were slight improvements in the number of complete and single-copy genes when the assemblies were upgraded to Hi-C, but this effect was minimal and only involved a few genes (Figure 1). Interestingly, all snow leopard genomes (even those upgraded with Hi-C or built with linked read technology) showed a larger number of missing genes, suggesting there may have been loss of conserved genes in that lineage during its evolutionary history. However, additional long-read sequencing should be incorporated to confirm this pattern.

**Figure 1:**
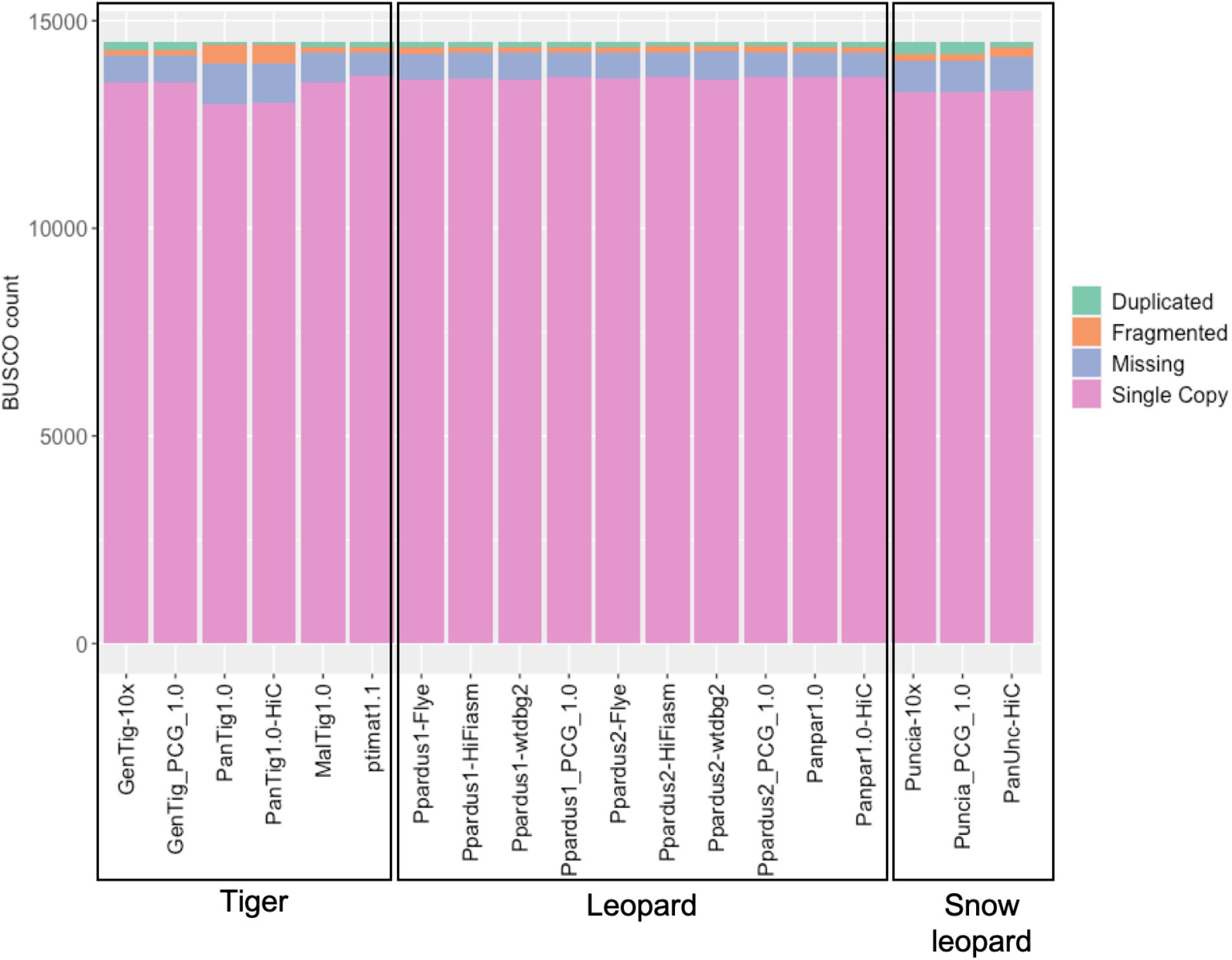
BUSCO scores from the mammalia_odb10 dataset across all draft (-), final (*), and previously published (unmarked) assemblies.

### Repeat content

We evaluated the repetitive content of each new assembly using a combination of homology and *de novo* repeat finding tools. Repeat content was consistent across the assemblies evaluated (Figure 2, 44.35%-45.51%) and this is mostly consistent with previous repeat content estimates for *Panthera [10]*. However, we did find that both of the long read leopard assemblies contained on average higher repetitive content compared to all other assemblies. Although we make no direct comparison here to a short read assembly from the same individuals, it does suggest that repeat discovery is improved by long read sequencing.

**Figure 2:**
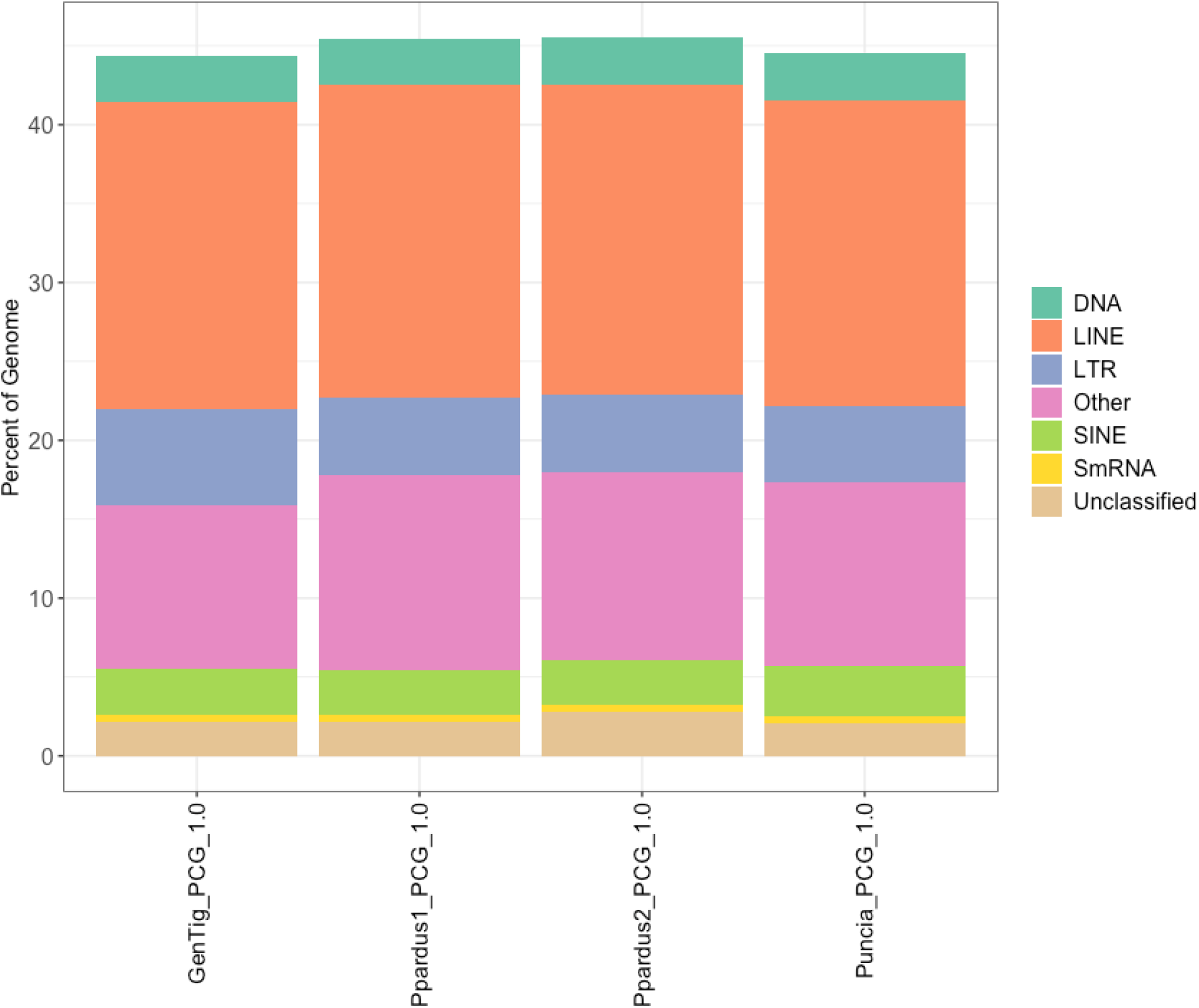
Percentages of various repetitive elements types for *de novo Panthera* genome assemblies.

### Whole genome alignment

We generated whole genome alignments for the domestic cat and the final *de novo* genomes for the leopard, snow leopard, and tiger. The whole genome alignments reiterate the conserved karyotype of the *Panthera* clade and additionally show the high continuity of each assembly (Figure 3). Additionally, we were able to assemble a majority of the X chromosome in each species. In the domestic cat, the X chromosome is estimated to be approximately 131.8Mb (GCA_000181335.5). In Ppardus1_PCG_1.0, we identified a scaffold corresponding to the X chromosome that was 126.8Mb in size, Ppardus2_PCG_1.0’s corresponding chromosome is 125.2Mb, GenTig_PCG_1.0’s putative X chromosome 120.3Mb, and lastly, Puncia_PCG_1.0 has a scaffold of 122.7Mb corresponding to the domestic cat X. This will enable further investigation into the peculiar lack of divergence of the X chromosome in the *Panthera* lineage, as reported by [8].

**Figure 3:**
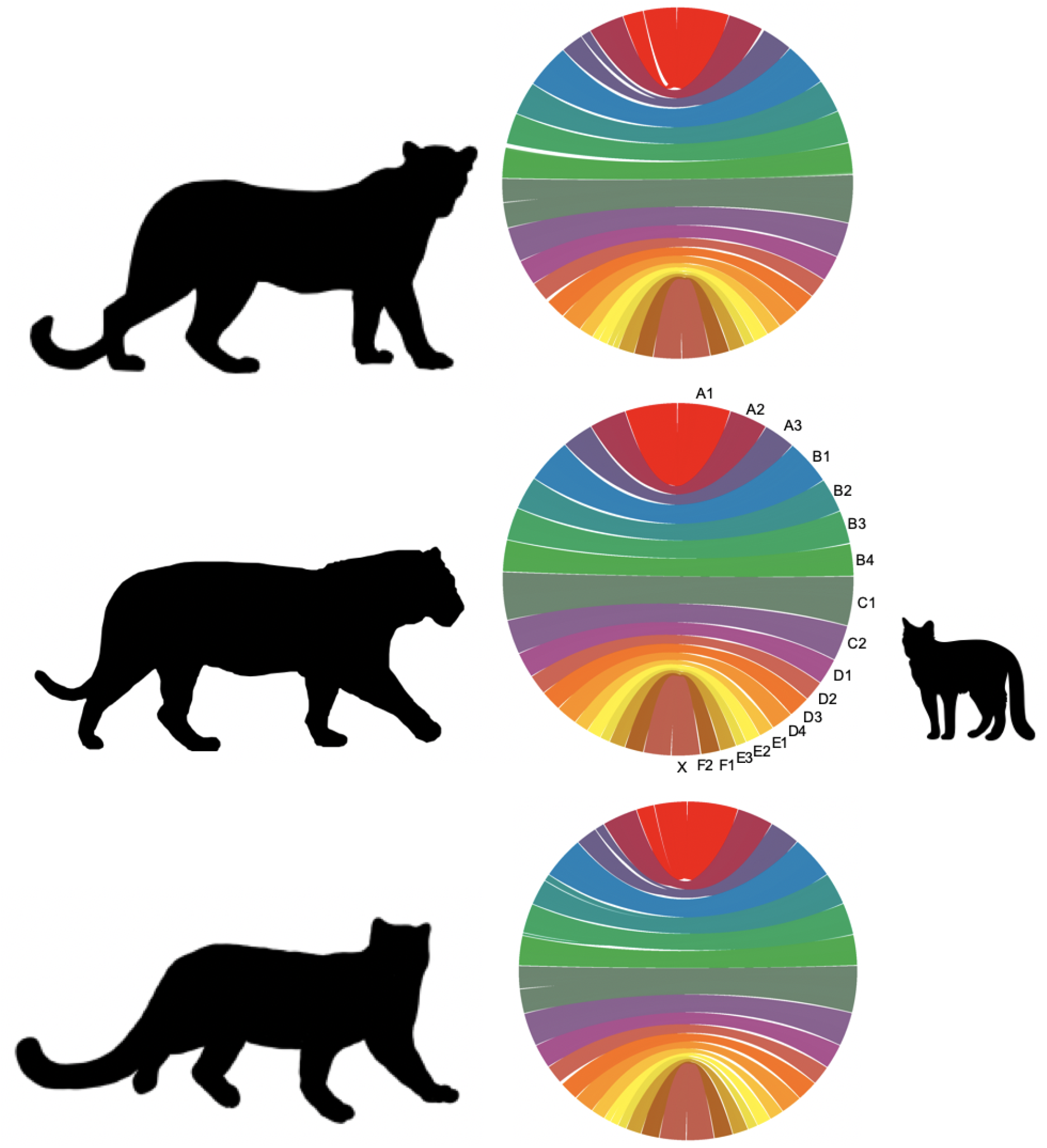
Whole genome alignments for (from top to bottom) leopard, tiger, and snow leopard (left) to the domestic cat (right). Silhouettes from phylopic.org: Snow leopard by Gabriela Palomo-Munoz ; leopard, no license; tiger by Sarah Werning; domestic cat, no license. http://creativecommons.org/licenses/by/3.0/.

### African leopard ancestry verification

Using principal component analysis, we analyzed 37 previously sequenced African leopards from [42] and the two leopards sequenced in this study with putative African ancestry. We found that these leopards fall well-within the bounds of what would be expected from an African leopard (Figure 4). An additional study analyzing multiple leopard subspecies [23] showed that African leopards are highly differentiated from every other leopard subspecies across their genomes, meaning we would expect that the leopards examined here would separate in PCA space if they were not African leopards. The two samples group most closely to leopards from Zambia, but additional data will be needed from across the African leopard range to identify the specific ancestry of these individuals. Although some additional African leopards were sequenced as a part of [23], most of these are from historic specimens. With continued land use change and habitat loss, a comprehensive modern examination of leopard structure across their range would assist in the identification of habitat corridors and further aid in confirming the source identification of poached individuals and other trafficked materials, such as the leopards here.

**Figure 4:**
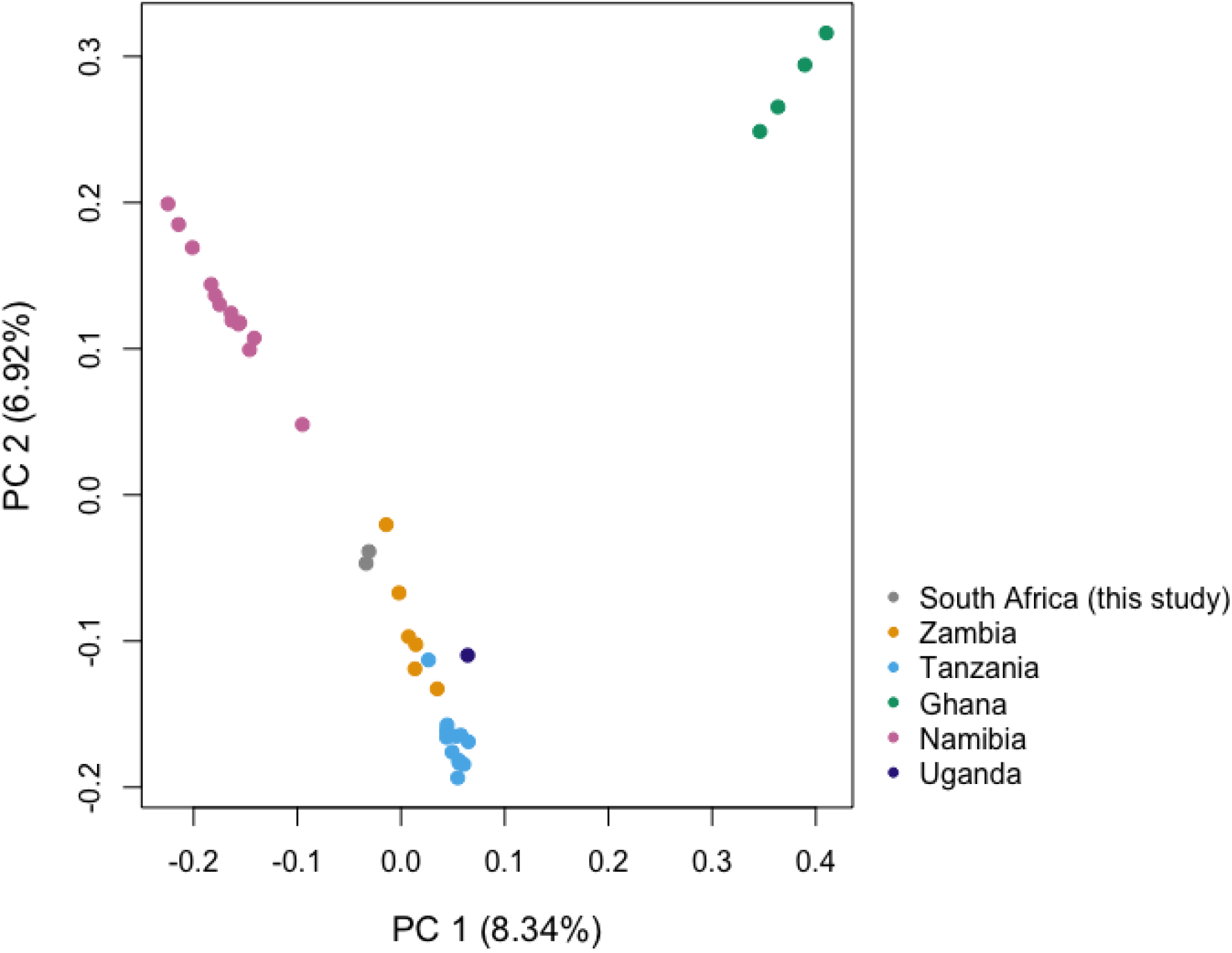
Principal component analysis showing PC1 vs PC2 of putative African leopards from this study, and previously published whole-genome data from African leopards. Samples are colored by source locations identified in [42].

### Phylogenetics

Using highly conserved genes from BUSCO, we built a tree of each (sub)species within the *Panthera* clade with available genomes, using the clouded leopard (*Neofelis nebulosa*) as the outgroup (Figure 5). The final tree was built using 8,719 single-copy orthologs. As has been previously reported, the quartet scores for each node indicate a substantial percentage of gene-tree discordance (42-67% of the gene trees supported each species-tree branch), but the overall species tree topology is supported with high bootstrap support (100). This is not completely unexpected, however, since we only included conserved coding regions. Our results also agree with previous studies that find the most discordance in the lion-leopard-jaguar trio [8,9]. More nuanced approaches using these high-quality genomes can be used to further investigate the phylogenetic discordance across the clade using large numbers of orthologous gene sequences as is done here or genome alignments that are not reliant on mapping to divergent species, as in previous studies [8,9].

**Figure 5:**
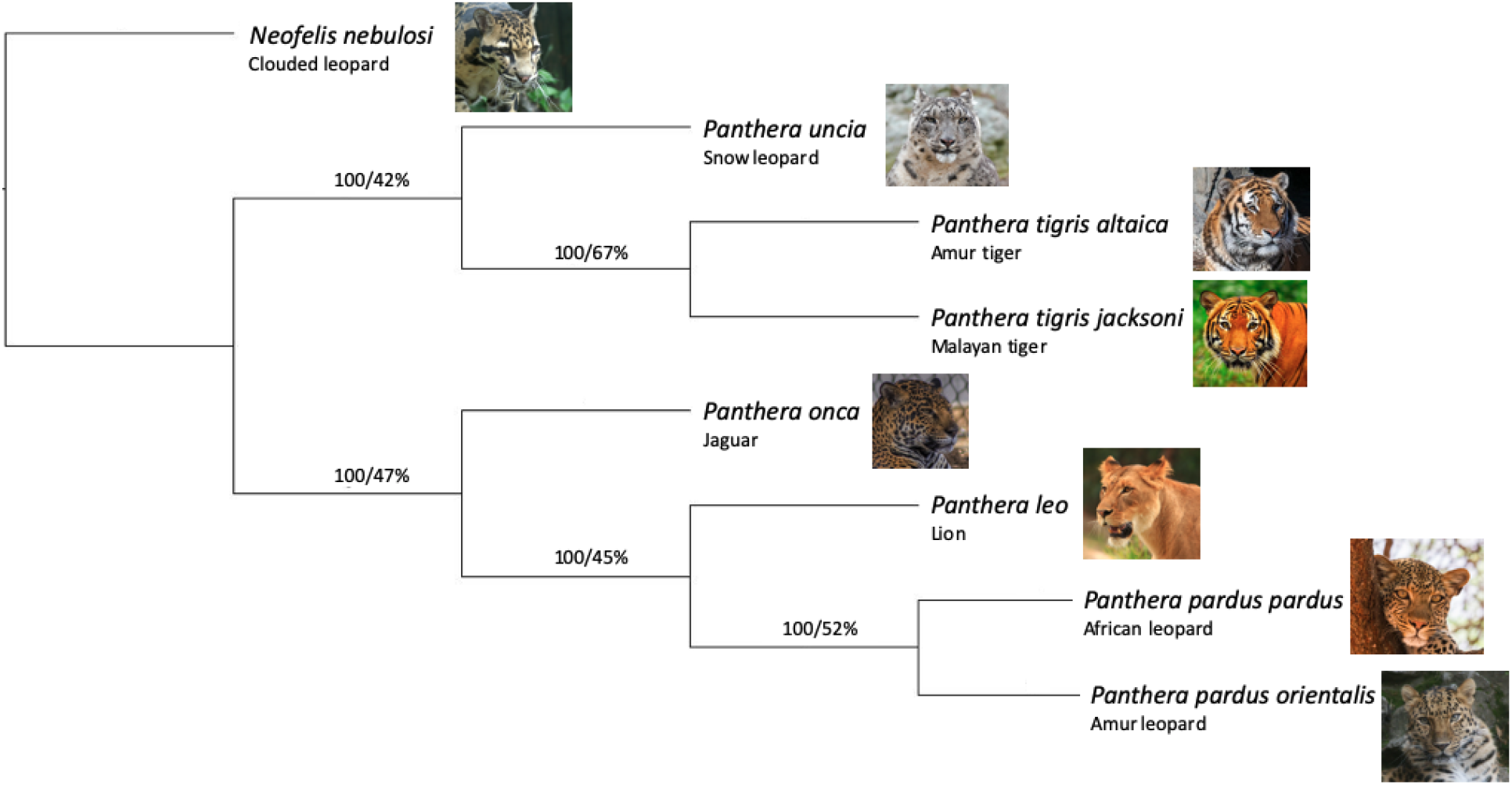
Cladogram representing phylogenetic results from ASTRAL-III using 8,719 BUSCO genes for the *Panthera* clade and an outgroup, the clouded leopard (*Neofelis nebulosa*). Plotted from results of the ASTRAL-III species tree using the best individual gene trees. Notation indicates bootstrap/quartet score for each node.

### Summary assessment of genome assembly and annotation

Using various technologies, we constructed several new chromosome-level assemblies for the tiger, the snow leopard, and the African leopard. We also use biobanked samples for PacBio sequencing and successfully built two high-quality assemblies from these samples. Our analysis showcases that older samples, subject to proper storage, can successfully be used for HMW DNA extraction and sequencing, reinforcing the utility and importance of biobanking initiatives.

Although several genomes exist for the tiger, we add a novel genome assembly for a captive tiger of unknown ancestry, an improved assembly for the snow leopard, and the first assemblies for the African leopard. We show that in all cases, incorporation of Hi-C data substantially increases the continuity of the genomes to approximately chromosome-level. Additionally, we find that as in previous studies, the *Panthera* clade is remarkably conserved at the karyotypic level, and find no substantial chromosome-level inversions between the domestic cat and any of the genomes examined. These high-quality reference genomes will help to reveal the evolutionary history of this important clade and are highly relevant to the conservation of *Panthera*.

## Data availability

All sequence data and final assembly files for the snow leopard have been deposited under NCBI BioProject PRJNA602938. All sequence data and final assembly files for the tiger have been deposited under NCBI BioProject PRJNA770127. All sequence data and final assembly files for the leopard have been deposited under NCBI BioProject PRJNA781109. Repeat annotation files available upon request.

## Acknowledgements

We wish to acknowledge In-Sync Exotics, specifically Vicky Keahey and Dr. Emily Wilson for assistance in obtaining tiger samples. In addition, we would like to thank Bill Nimmo and Kizmin Reeves of Tigers in America for facilitating access to the tiger sample. We would like to thank Jenny Brubaker, RVT and Jennifer Sohl, RVT of Maryland Zoo for granting us access to leopard samples and for sharing their stories of leopards Hobbes and Amari. Leopard sequencing was made possibly by a PacBio SMRT sequencing grant awarded to E.E. Armstrong and we are grateful to Pacific Biosciences, specifically Jay Morris, for assistance in the leopard genome sequencing and assembly. We would also like to thank the Georgia Genomics and

Bioinformatics Core for their assistance in HiFi sequencing. We would like to thank the San Diego Frozen Zoo for access to the snow leopard sample. We also acknowledge Dhananjay Wagh at the Stanford Functional Genomics Facility for his assistance with high molecular weight extraction and library preparation of the snow leopard sample. Figure licenses for Figure 5: “Malayan tiger” by zambase, “Clouded Leopard” by tim ellis, “Snow Leopard Relaxed” by Eric Kilby, “Jaguar” by Princess Stand in the Rain, “African Leopard - Panthera pardus pardus, Tsavo West NP, Oct 13-5” by Peter R Steward, and “Amur Leopard” by Tasshu Rikimara. Amur tiger is CC0.

